# Microbial Degradation of Contaminants of Emerging Concern by Pseudomonas putida in Wastewater Treatment Plants

**DOI:** 10.1101/2024.09.23.614627

**Authors:** Boyu Lyu, Bharat Manna, Xueyang Zhou, Ivanhoe K. H. Leung, Naresh Singhal

## Abstract

Contaminants of Emerging Concern (CECs) are a class of contaminants that are commonly found in urban wastewater treatment plants (WWTP) at low concentrations. These compounds include antibiotic drugs, personal care products, and industrial chemicals, which are widely used in modern society. Although they provide undeniable benefits in our daily lives, their accumulation in the environment poses a significant risk to human health and the ecosystem. Due to the incomplete removal of these compounds by traditional WWTPs, there is a need to understand the interactions between microbes and CECs to develop effective solutions to this environmental challenge. *Pseudomonas putida* is a bacterium commonly found in water-related habitats, including freshwater streams, forests, and WWTPs. It is known for its ability to degrade various organic compounds, making it a suitable model for understanding the linkages between bacterial enzymes and CECs found in WWTPs. In this study, isolated *Pseudomonas putida* strain KT2440 was cultured in lab-scale reactors to mimic the WWTP environment. The degradation of this isolated strain towards different groups of CECs were studied over a 24-hour period using liquid chromatography with tandem mass spectrometry (LC-MS/MS). Mass spectrometry-based quantitative proteomics was also performed at different sampling points to establish a linkage between the enzyme expression profile of this strain and its degradation of CECs via bioinformatics analysis. The results of this study demonstrated that *Pseudomonas putida* KT2440 is capable of degrading atrazine, acetamiprid, carbendazim, diclofenac acid, erythromycin, and sulfamethazine at lab scale. At the 20-hour time point, the enzyme expression profile indicated a high protein abundance of various oxidoreductases that could explain the observed CEC degradation.The study’s findings provide important insights into the potential of Pseudomonas putida as a bioremediation agent for CECs in WWTPs. The degradation of CECs by *Pseudomonas putida* KT2440 is a significant contribution to the bioremediation process, and its enzyme expression profile can serve as a case study for more complex interactions of microbes and CECs in real WWTPs. Furthermore, this study supports the potential of *Pseudomonas putida* as a promising strain for various bioremediation applications. Finally, this study’s results can help develop more efficient and effective methods for removing CECs from wastewater, which can contribute to the protection of public health and the environment.

## 1.0 INTRODUCTION

Contaminants of Emerging Concern (CECs), including pharmaceutical residues, are increasingly recognized as a significant environmental concern, particularly due to their persistence in wastewater (Huang et al., 2020). Conventional wastewater treatment processes, such as activated sludge systems, often fail to fully degrade these compounds, leading to their accumulation in aquatic ecosystems (Brunelle et al., 2024). This accumulation poses substantial ecological threats, as CECs can traverse the food chain and reach threshold concentrations that may induce reproductive and developmental issues in aquatic organisms (Fu et al., 2021). Such impacts raise alarming implications for human health, especially considering the growing scarcity of clean drinking water in many regions worldwide. The contamination of water resources with CECs endangers biodiversity and threatens the health and well-being of communities reliant on these resources (Moreira et al., 2016).

In contrast to urban areas equipped with advanced wastewater treatment infrastructures, rural regions—particularly those with extensive agricultural activities and aquaculture—face unique challenges (Ahmed et al., 2017). These areas often utilize significant quantities of antibiotics for disease control, leading to substantial pollution of local water bodies (Ahmed et al., 2017). The lack of effective treatment options and primary processing units to manage local wastewater exacerbates the situation, presenting even more severe challenges to vulnerable communities that may lack the resources to address these contaminants effectively (Ahmed et al., 2017). While biological treatment processes remain among the most cost-effective methods for treating wastewater, traditional approaches struggle to address the emerging threat posed by CECs. Microbial communities, composed of millions of diverse organisms, play a crucial role in the biological treatment of wastewater (Oberoi et al., 2019). Each member of these communities contributes unique metabolic capabilities, particularly in terms of carbon conversion and nutrient removal (Ren et al., 2018). Different bacterial species exhibit varying abilities to remove CECs; those that can thrive in diverse environmental conditions and utilize a wide range of nutrient sources are often better suited for CEC degradation. Notably, these microorganisms harbor essential degradation enzymes, including oxidoreductases, hydrolases, and, most importantly, microbial cytochrome P450s, which are particularly adept at degrading complex chemical structures (Ahmed et al., 2017).

Among the identified *gammaproteobacteria, Pseudomonas putida* stands out for its remarkable versatility in metabolizing a wide array of organic compounds, positioning it as a promising candidate for bioremediation applications (Haernvall et al., 2017). With a size of just a few micrometers, this microorganism can effectively function as a CEC degrader in low-cost wastewater treatment units, avoiding the need for heavy investments in advanced technologies (Xiao et al., 2018). However, the full extent of this strain’s ability to degrade CECs remains underexplored, particularly concerning how it adapts to varying oxygen availability in low-end wastewater treatment systems that lack adequate aeration. Understanding these dynamics is crucial for leveraging the potential of *Pseudomonas putida* in wastewater treatment processes (Haernvall et al., 2017).

This study aims to investigate the degradation capabilities of *Pseudomonas putida* in lab-scale reactors designed to mimic different dissolved oxygen levels found in real-world decentralized wastewater treatment units (Oberoi et al., 2019). By assessing its efficiency in degrading a range of CECs in synthetic wastewater, this research seeks to provide insights into the practical application of *Pseudomonas putida* as an effective CEC degrader. Connecting to proteomics, we have insight into how microbial enzymes could cope with CECs for degradation. Ultimately, the findings from this study could pave the way for implementing cost-effective biological treatment processes that benefit local communities, enhancing their ability to manage wastewater and protect vital water resources.

## 2.0 MATERIALS AND METHODS

### 2.1 REACTOR OPERATION

The experiment was conducted using six identical 1L bioreactors that were operated simultaneously at a controlled room temperature of 20 ± 1°C. Over a 24-hour incubation period, each bioreactor experienced distinct aeration conditions (Figure 1&Figure S1). In condition A, the dissolved oxygen (DO) concentration was kept constant at 2.0 ± 0.2 mg/L. Condition B involved a dynamic DO level that fluctuated in 3-minute cycles, where the DO rose from 0 to 2.0 ± 0.2 mg/L and then fell back to 0 mg/L. Condition C featured cyclic DO changes within 6-minute cycles; during each cycle, the DO level increased from 0 to 2.0 ± 0.2 mg/L over 3 minutes, followed by a reduction back to 0 mg/L, after which the DO was maintained at 0 mg/L for another 3 minutes. Conditions D, E, and F mirrored the operational strategies of conditions A, B, and C, respectively, but utilized a different DO range of 0 to 8 mg/L.

**Figure 1.**
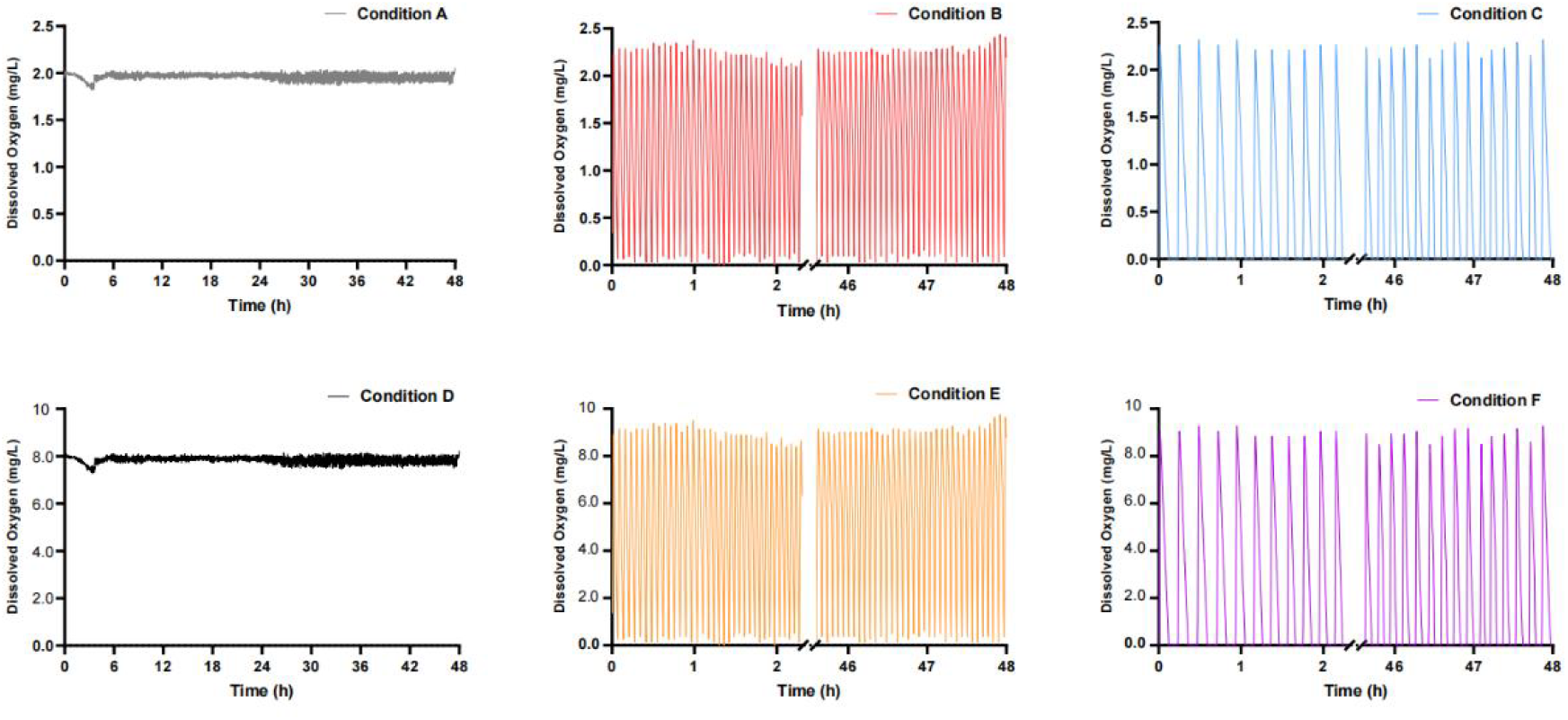
Reactor operations DO ranges, A Constant aeration at 2mg/L (CA), B Continuous perturbation (CP) at DO of 2mg/L, C Intermediate perturbation (IP); Conditions D, E, F are same strategy but at DO from 0∼8mg/L.

The three scenarios that implemented fixed air flow rates of 0.5 L/min utilized a mass flow controller and compressed air to achieve rapid increases in DO. Each reactor was initially added precultured *Pseudomonas putida* KT2440 at an optical density of OD 0.1 and 1 mL of a trace element solution. A synthetic wastewater solution (Table 1) was continuously supplied to the reactors through a syringe pump and thoroughly mixed using a magnetic stirrer.

**Table 1.** Trace elements and synthetic wastewater provided to the reactor.

| Chemical | Concentration (g/L) |
| --- | --- |
| EDTA | 2.50 |
| ZnSO<sub>4</sub>·7H<sub>2</sub>O | 1.10 |
| CoCl<sub>2</sub>·6H<sub>2</sub>O | 0.80 |
| MnCl<sub>2</sub>·4H<sub>2</sub>O | 2.55 |
| MgSO<sub>4</sub>·7H<sub>2</sub>O | 20.0 |
| CuSO<sub>4</sub>·5H<sub>2</sub>O | 0.86 |
| (NH<sub>4</sub>)<sub>6</sub>Mo<sub>7</sub>O<sub>24</sub>·4H<sub>2</sub>O | 0.07 |
| CaCl<sub>2</sub>·2H<sub>2</sub>O | 2.75 |
| FeSO<sub>4</sub>·7H<sub>2</sub>O | 2.57 |
| Synthetic wastewater | Concentration |
| Methanol (mL/L) | 26.94 |
| NH<sub>4</sub>Cl (g/L) | 24.46 (C); 30.57 (B & A) |
| KH<sub>2</sub>PO<sub>4</sub> (g/L) | 2.76 |
| K<sub>2</sub>HPO<sub>4</sub> (g/L) | 2.76 |
| NaHCO<sub>3</sub> (g/L) | 76.80 |

### 2.2 CONTAMINANTS OF EMERGING CONCERN ANALYSIS

CECs were purchased from AK Scientific, USA, with the highest analytical purity, and stock solutions were made for CECs at 20 ppm with methanol (Merck; LC-MS grade purity). The working solution was prepared prior to the experiment, and the 1L reactor was spiked with 10μg/L of the 24 CECs. To extract the remaining CECs, solid phase extraction (SPE) was performed using OASIS Prime HLB cartridges (Waters, Milford, MA, U.S.A.) with a standard method. At the end of the cultivation period, 200 mL of the bioreactor culture was collected and centrifuged (10,000g, 4 °C, 20 min) to remove suspended particles and cell pellets. The supernatant was acidified to pH=2 using 0.1M HCl and then passed through cartridges, washed with 5% methanol, and finally, the CECs were eluted with 100% methanol. The extracted CECs were analyzed using a Shimadzu 8040 LC-MS/MS instrument (Shimadzu, Japan) with a C18 column (2.1 × 100 mm2, particle size 1.8 μm, Agilent Technologies, Germany). A binary gradient of 0.1% formic acid in deionized water (mobile phase A) and 0.1% formic acid in methanol (mobile phase B) were used for both ESI + and ESI - model. The flow rate for both modes was 0.25 mL/min for 20 mins, and internal standards were used to minimize matrix effects. Limits of detection and quantification were set with signal-to-noise ratios 10, and OMP removal efficiency was calculated for 48 hours via Prism 9, respectively.

### 2.3 ENZYME IDENTIFICATION

Proteomic analysis was performed on 5 mL sludge samples under various experimental conditions, with three biological replicates per condition. Sludge samples were pelleted and washed, and protein extraction was conducted by sonication and centrifugation. Purification was achieved with SpeedBead carboxylate-modified E3 and E7 magnetic particles (Sera-Mag, USA) to yield 150μg protein for each sample with EZQ protein assay kit to quantify the protein concentration. The protein extract was then reduced with 5 mM dithiothreitol (DTT), alkylated with 15 mM iodoacetamide, and digested with 1.5 µg trypsin. Peptides were cleaned by solid-phase extraction (Oasis Prime HLB 1cc, 30 mg), and concentrated using a speed vacuum. Peptide samples (10 µL) were analyzed by nano LC-MS/MS using a NanoLC 400 UPLC system and a TripleTOF 6600 Quadrupole-Time-of-Flight mass spectrometer. Data were searched against a metagenomics-derived database using MetaProteomeAnalyzer and X-tandem, with a 1% false discovery rate filter. The resulting group file was converted to MetaProteomeAnalyzer for metaproteomics analysis and clustering. Enzyme regulation information was obtained from RegulonDB v12.0. The enzyme CECs correlation was achieved with mental test in R package (Ver 4.3.3)

## 3.0 RESULTS AND DISCUSSION

### Degradation of CECs by *Pseudomonas Putida* KT2440

In this study, the degradation of atrazine, acetamiprid, carbendazim, diclofenac acid, erythromycin, and sulfamethazine by *Pseudomonas putida* was evaluated over 24 hours under six different aeration conditions. The results, depicted in Figure 2&Figure S2 show significant differences in the residual percentage of the compounds depending on the aeration levels. For instance, atrazine under low aeration conditions (IP8) exhibited between 82% and 61% residuals across three replicate reactor runs. Other aeration conditions incorporate less anoxic phases, resulting in significantly better degradation of atrazine to the lowest residual of 45% under the CP2 condition. In comparison, the duration of the aeration period in batch reactors significantly influences this compound’s removal efficiency, and the degradation can be optimized by supplying sufficient oxygen into the system. As CA2 conditions have constant aeration of DO at 2 mg/L and IP2 processes the lowest average DO of 0.5 mg/L, our study demonstrated the optimized aeration strategy to reduce 75% of the need for total DO supply and retain the relative removal for these CECs (Achermann et al., 2018).

**Figure 2.**
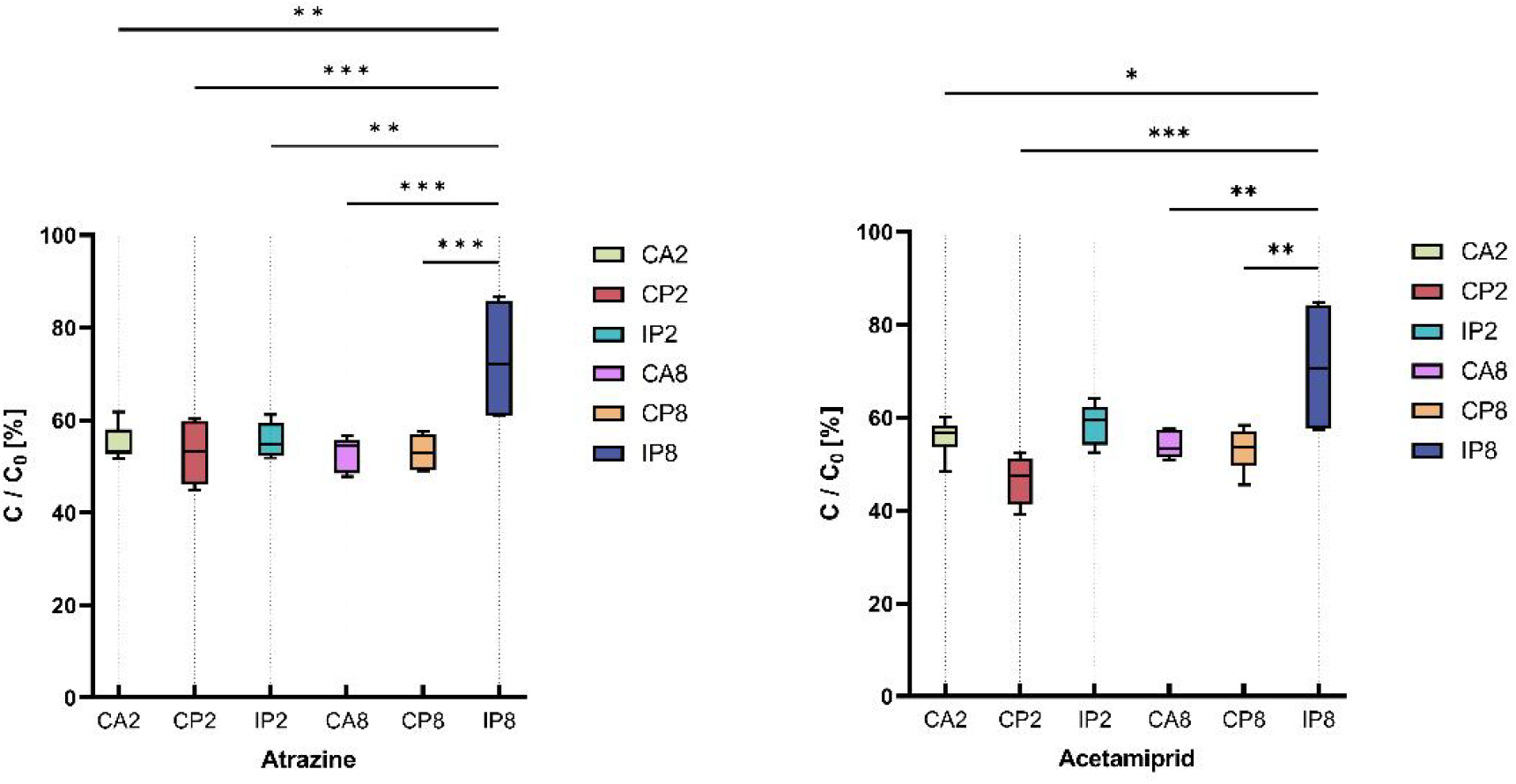
Removal of important CECs of atrazine and acetamiprid by Pseudomonas putida in lab scales under different aeration conditions.

Comparable degradation results have been identified for other CECs, such as acetamiprid. In Figure 2, the IP8 condition showed the highest residual of over 60%, while other conditions, such as CP2, reduced the CEC residual to as low as 40%, suggesting a relative 20% increase in degradation if switched to the optimum aeration conditions. Unlike atrazine, the CP2 condition tends to have better removal capability over the other conditions, with an average DO level of 1 mg/L. This could be due to the different metabolism rates of *Pseudomonas putida* KT-2440 under distinct aeration conditions and the differences in enzyme expression profiles. The degradation of atrazine and acetamiprid is of significant environmental importance due to these compounds’ persistent and harmful nature. Atrazine, a widely used herbicide, is known for its persistence in soil and water, posing risks to aquatic life and potentially contaminating drinking water sources. Acetamiprid, a neonicotinoid insecticide, has potential impacts on human health through food residues. It is important to acknowledge the threat of CECs and mitigate their discharge into the environment.

Figure 3 demonstrates the removal of other CECs like carbendazim, a fungicide that is persistent in the environment and can remain in soil and water for extended periods. The CEC-degrading strain *Pseudomonas putida* KT-2440 in this study has demonstrated relatively good removal, with the lowest residual observed being 20% after 24 hours of reactor incubation. The concentration of carbendazim was reduced from 10 μg/L to close to 2 mg/L, achieving an average removal efficiency of 80%. This suggests a promising solution for the bioremediation of carbendazim in wastewater, given its toxicity to a wide range of aquatic organisms, including fish and invertebrates (Shao et al., 2019). No significant residual difference was observed among the tested aeration conditions, indicating that DO is not a limiting factor and that WWTPs are not. Diclofenac acid, on the other hand, shows a significant dependence on the DO level present in this system. As indicated in Figure 3, the lowest residual observed was 47% under the CP2 condition, while the other aeration conditions limited the degradation to less than 40%. Interestingly, the CA2 condition did not yield the best removal efficiency. We hypothesize that enzyme expression under CP2 plays a critical role in degrading this CEC. Notably, the accumulation of diclofenac acid poses a threat to aquatic life, especially fish, and can lead to higher concentrations up the food chain, potentially impacting predators, including birds and mammals (Kanaujiya et al., 2019).

**Figure 3.**
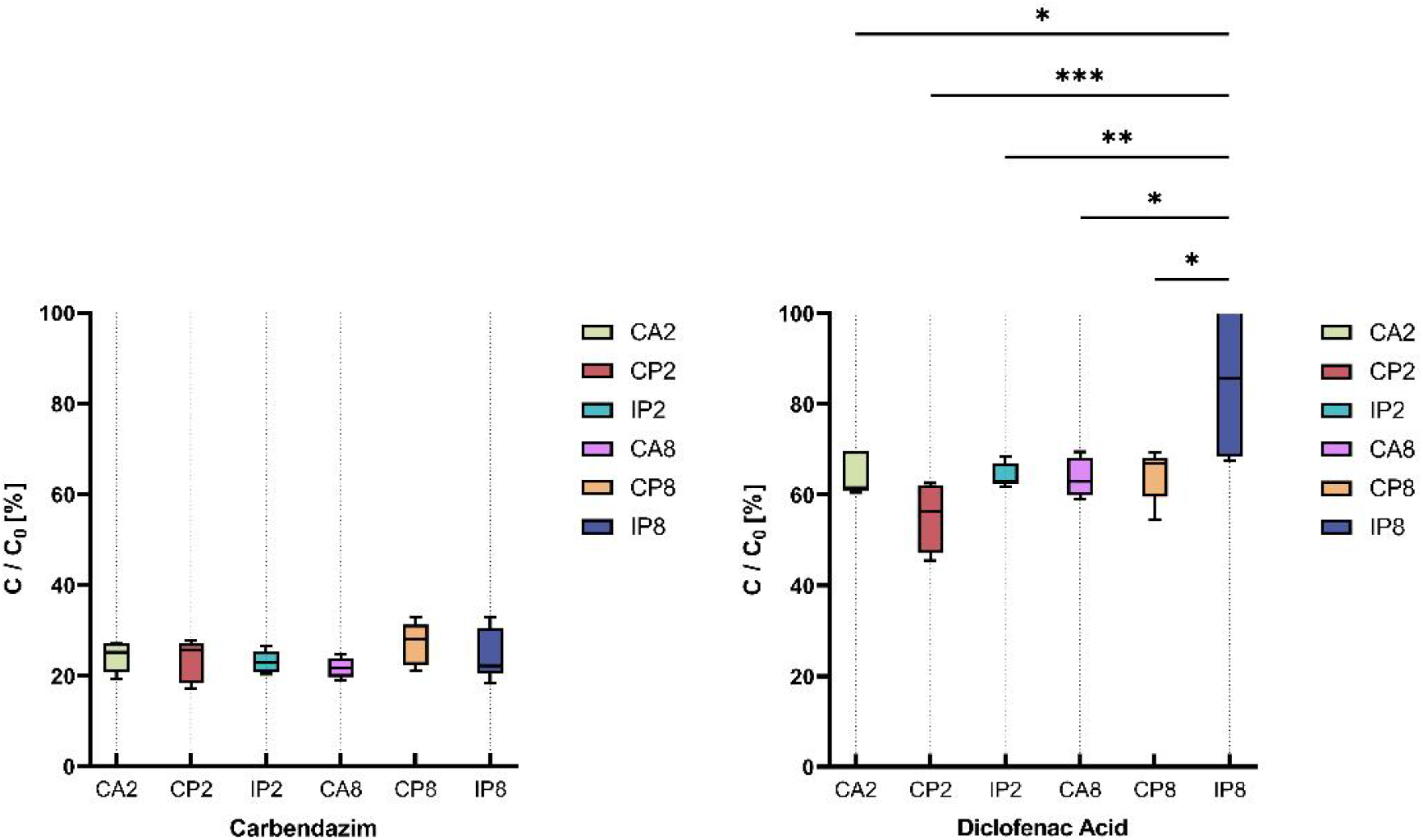
Removal of important CECs of carbendazim and diclofenac acid by Pseudomonas putida in lab scales under different aeration conditions.

Finally, with the CEC-degrading strain *Pseudomonas putida* KT-2440, we observed the successful removal of the antibiotic erythromycin, achieving over 90% removal on average across different aeration conditions. As shown in Figure 4, after 24 hours of incubation, the majority of erythromycin was biodegraded, with no significant difference among conditions. This CEC is known to promote antibiotic resistance with environmental exposures. We encourage future studies to investigate the degradation products of this CEC since over 90% of the parent erythromycin has been transformed, and to assess the associated toxicity (Shao et al., 2019). Sulfamethazine is another persistent antibiotic drug residue in the environment, and its removal is vital to mitigate the threat of potential resistant strains. Our data in Figure 4 show significant improvements in this CEC under both CP2 and IP2 conditions compared to CA2, which has a constant aeration DO of 2 mg/L. The lowest residual observed in batch reactors with the CEC-degrading strain *Pseudomonas putida* KT-2440 reached 23%, indicating that a reduced DO supply could promote the removal of this and presumably other compounds with similar structures. Importantly, we hypothesize that the enzyme expression profile of *Pseudomonas putida* KT-2440 under prolonged anoxic phases (IP2 and IP8) is critical for the observed removal differences (Shao et al., 2019).

**Figure 4.**
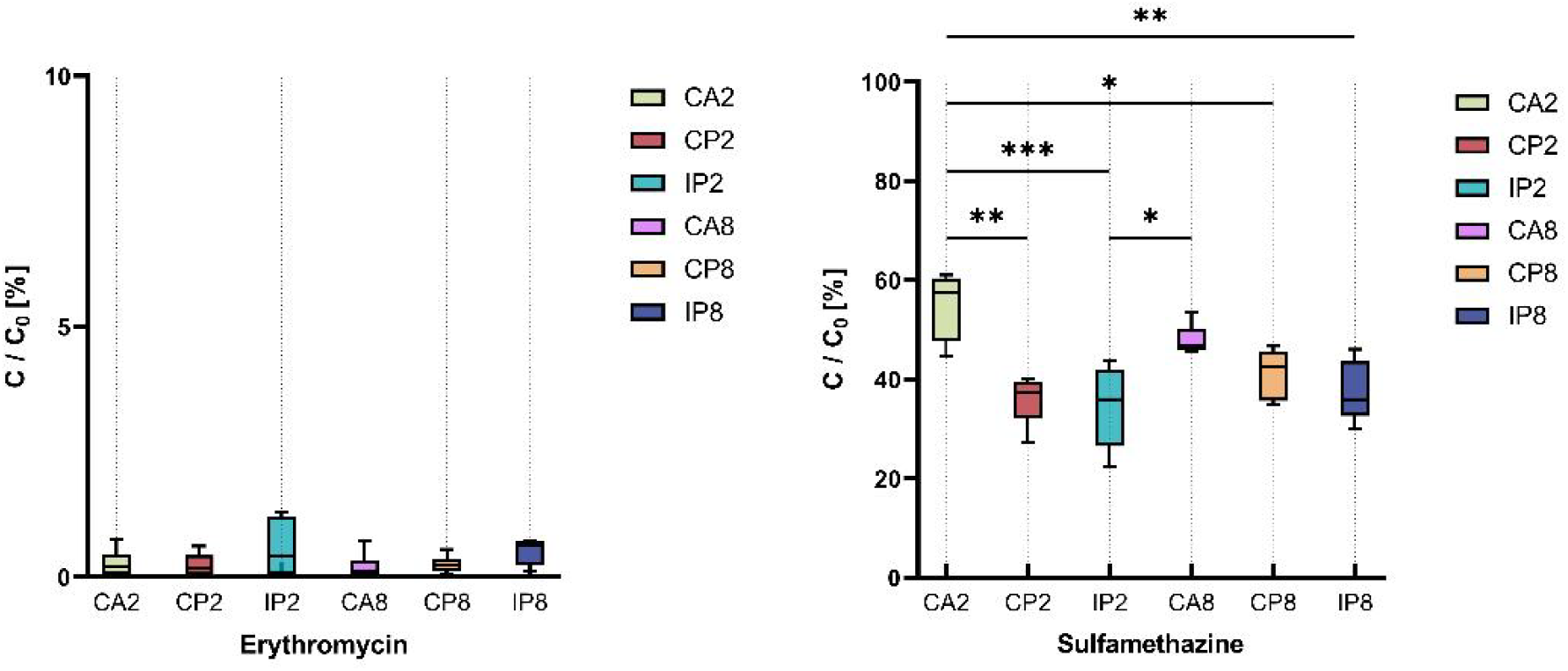
Removal of important CECs of erythromycin, and sulfamethazine by Pseudomonas putida in lab scales under different aeration conditions.

### Enzyme Expression Profiles of *Pseudomonas Putida* KT2440

In order to understand the distinct degradation behavior of the CEC-degrading strain *Pseudomonas putida* KT-2440 and optimize its degradation for CECs, we applied proteomic analysis to review enzyme expression under different conditions and its correlation to individual compounds. The data shows 27 unique oxidases among the tested conditions, and their distribution is illustrated in Figure 5. Oxidase is one of the most important groups of microbial enzymes for CEC degradation, and we observed significant correlations between detected oxidases and the removal of individual CECs. We identified a relatively higher presence of amine oxidase in the CP2 condition throughout the 24 hours of incubation compared to the CA2 condition, which has a constant 2 mg/L DO.

**Figure 5.**
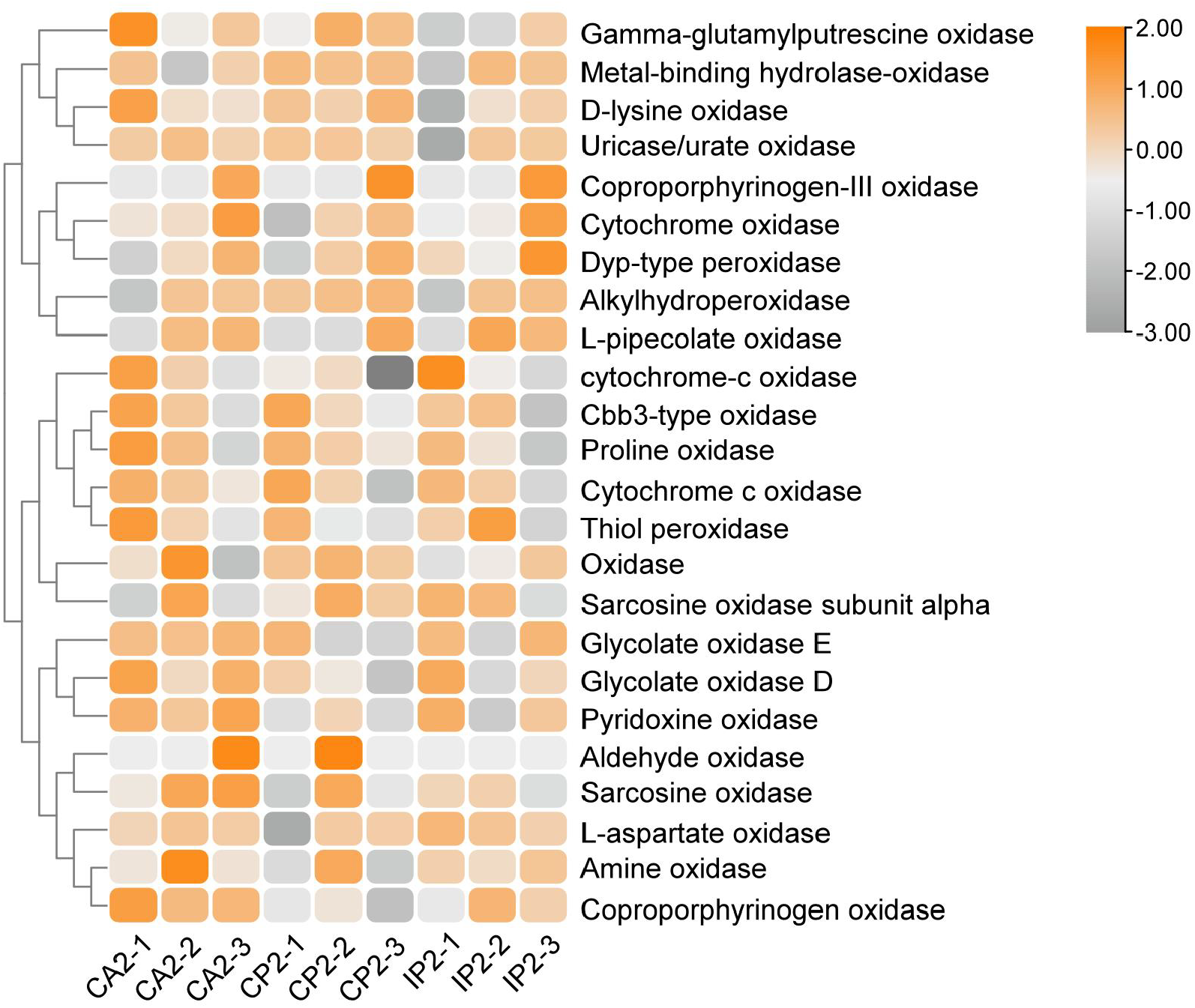
Protein expression under different aeration conditions for Pseudomonas Putida KT2440, normalized abundance from 2 to -3.

The changes in amine oxidase could be attributed to metabolic switches when an additional anoxic period was introduced to the bacteria. Typically, amine oxidase helps bacteria utilize organic amine-containing compounds in the environment as their nitrogen source, increasing their chances of adaptation and survival. *Pseudomonas putida* KT-2440 has been found to cope with diverse environments like lakes, soil, and wastewater. The insufficient oxygen supply in conditions like CP2 or IP2 intentionally stimulates the bacteria. The expressed amine oxidase would then oxidize and degrade various amine-containing CECs.

Other oxidases shown in Figure 6 have demonstrated their individual linkage to different CECs through statistical tests, indicating they could be additional CEC degraders. However, it is important to acknowledge that statistical correlation does not necessarily imply causality. We encourage future studies to confirm the reaction mechanisms via pure enzyme assays. For application prospects, we are more focused on the removal of CECs.

**Figure 6.**
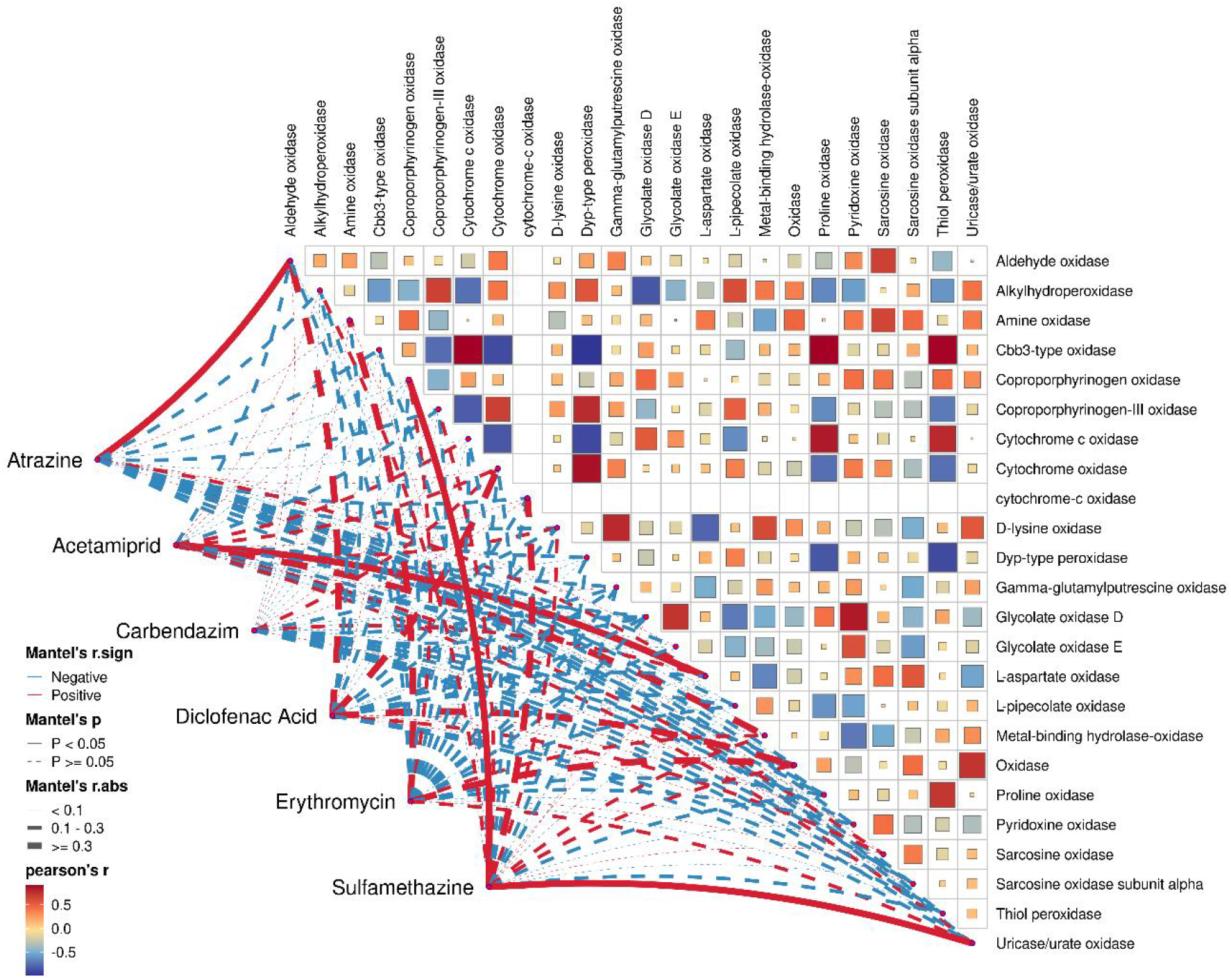
Mental analysis of expressed oxidases and different of CECs for correlation analysis.

## 4.0 CONCLUSIONS

The findings of this study highlight the significant role of *Pseudomonas putida KT2440* in the degradation of various CECs under different aeration conditions. The degradation efficiency of atrazine, acetamiprid, carbendazim, diclofenac acid, erythromycin, and sulfamethazine by *Pseudomonas putida* was influenced by the oxygen levels in the reactors. Specifically, conditions with dynamic aeration, such as CP2 (continuous perturbation at DO of 2 mg/L), generally resulted in higher degradation efficiencies compared to constant aeration conditions like CA2.The proteomic analysis revealed that the expression of certain oxidoreductases, particularly amine oxidase, was higher under dynamic aeration conditions, correlating with improved degradation of amine-containing CECs. This suggests that *Pseudomonas putida adapts* its metabolic pathways in response to fluctuating oxygen levels, thereby enhancing its degradation capabilities.

The practical application of these findings in WWTPs could revolutionize the current approaches to handling CECs. Traditional WWTPs often struggle with the complete removal of CECs, leading to their accumulation in the environment and posing risks to both ecosystems and human health. The ability of *Pseudomonas putida KT2440* to effectively degrade CECs under varying aeration conditions presents a promising solution (Haernvall et al., 2017). By incorporating dynamic aeration strategies similar to the CP2 condition used in this study, WWTPs can optimize the degradation of CECs. This approach not only improves the removal efficiency of persistent pollutants like atrazine and acetamiprid but also reduces the need for constant high oxygen supply, making the process more energy-efficient and cost-effective. The study’s findings are particularly relevant for decentralized wastewater treatment units, which often lack advanced aeration infrastructure. Implementing variable aeration regimes could enhance the degradation performance of microbial communities in these systems, thereby improving water quality in rural and underserved regions (Haernvall et al., 2017).

In conclusion, the study underscores the potential of *Pseudomonas putida* KT2440 as a robust bioremediation agent in WWTPs. By adopting optimized aeration strategies and leveraging the bacterium’s enzymatic capabilities, it is possible to achieve more efficient and sustainable treatment of wastewater, ultimately contributing to the protection of public health and the environment. Future research should focus on scaling these findings to pilot and full-scale WWTPs, as well as exploring the long-term stability and adaptability of *Pseudomonas putida* in diverse operational settings.

## ACKNOWLEDGMENT

We thank New Zealand eScience Infrastructure (NeSI) for computing support. Funding Sources Supported by the Marsden Fund Council, New Zealand Royal Society Te Apārang. [grant number MFP-UOA2018].

