## Supplemental Data for "Microbial Degradation of Contaminants of Emerging Concern by Pseudomonas putida in Wastewater Treatment Plants"

**Supplementary Figures**

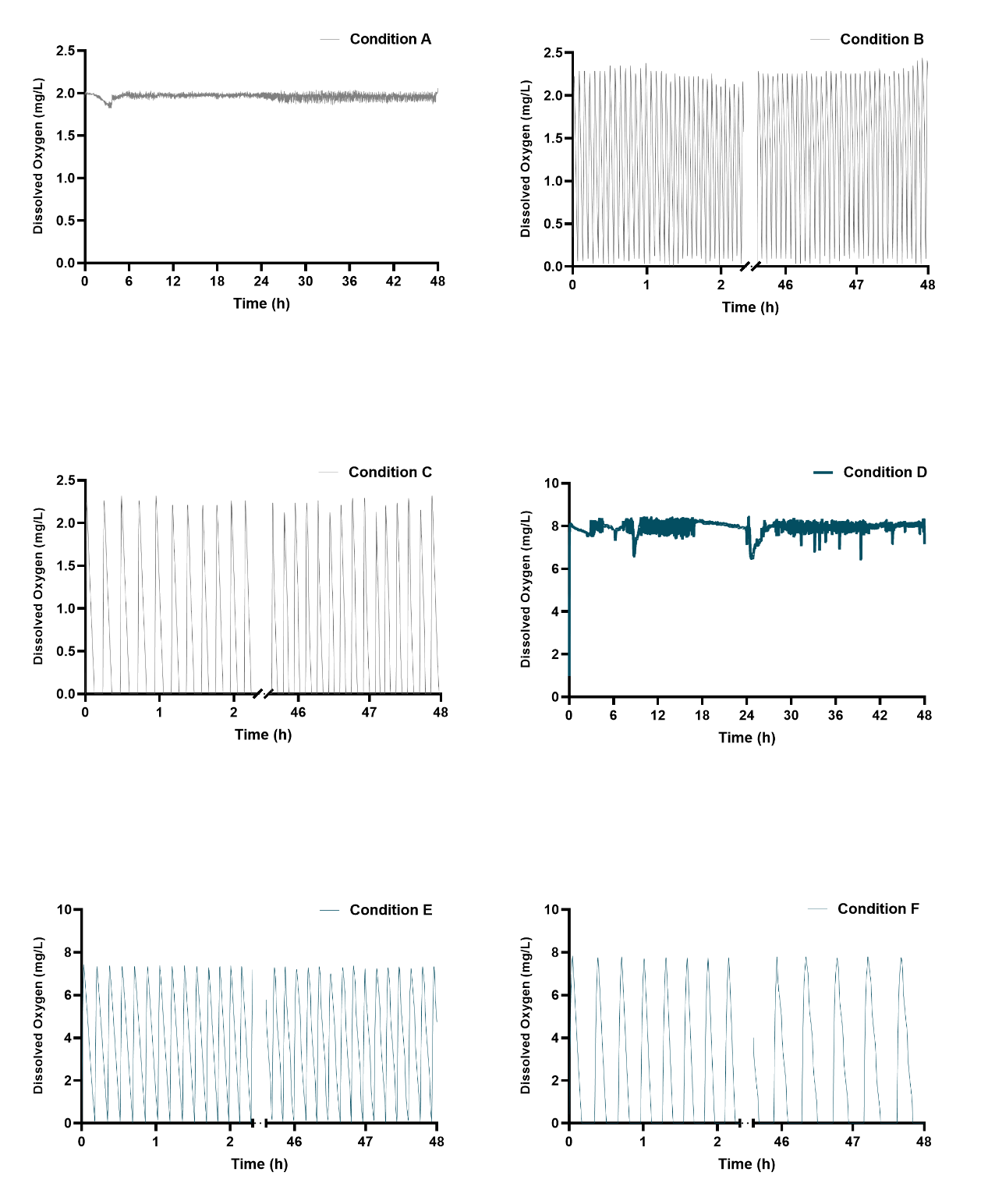

**Figure S1** Dissolved Oxygen Concentration of Conditions A, B, and C（Exp 1.）and D, E, and F (Exp 2.)

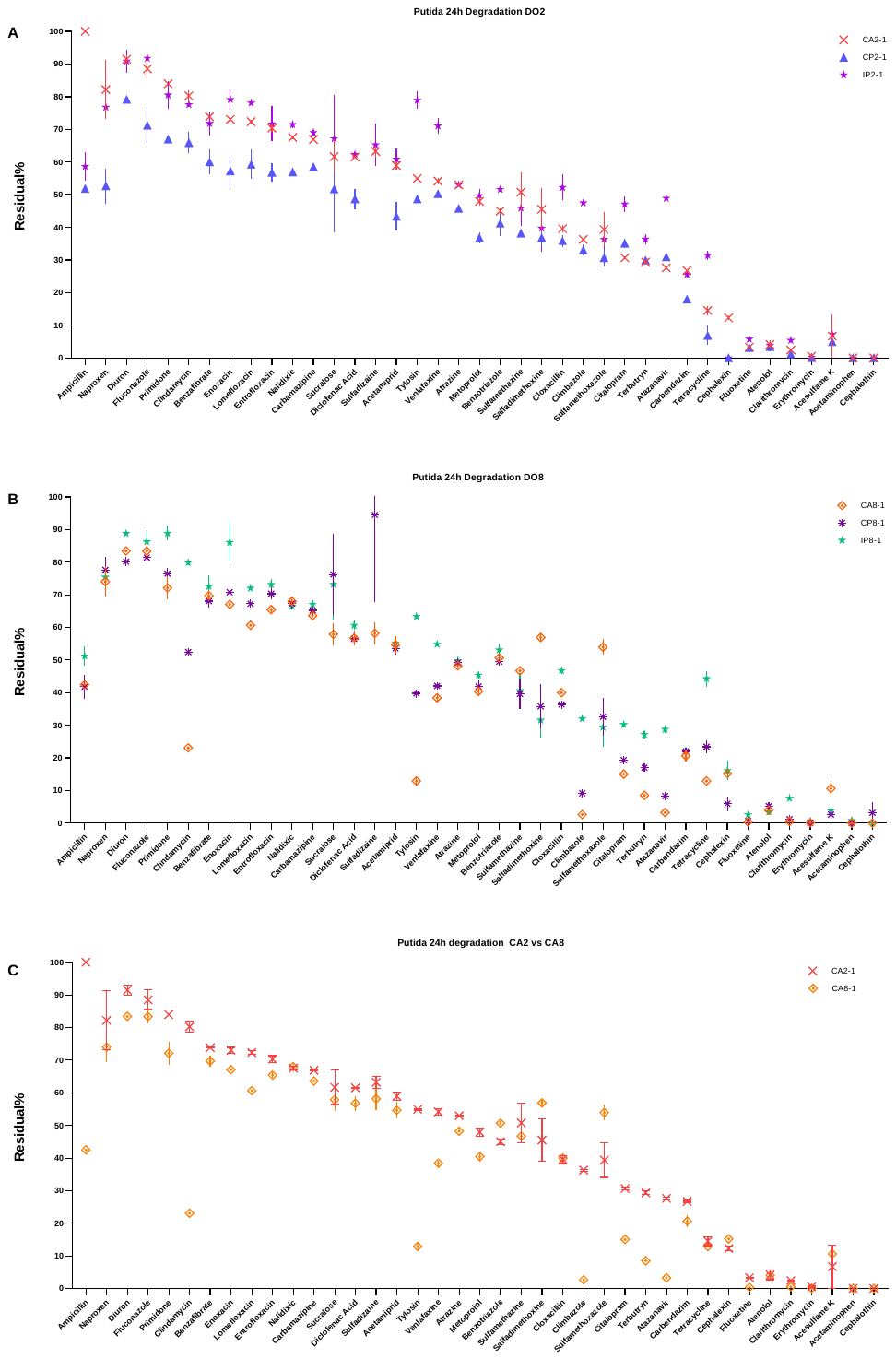

**Figure S2.** *A). Pseudomonas Putida OMP degradation residual at 24 hours under CA2, CP2, and IP2 conditions. (B) Residual of OMPs under CA8, CP8, and IP8 conditions. (c) OMP residual under CA2 and CA8 conditions.*
